# Euglenozoan kleptoplasty illuminates the early evolution of photoendosymbiosis

**DOI:** 10.1101/2022.11.29.517283

**Authors:** Anna Karnkowska, Naoji Yubuki, Moe Maruyama, Aika Yamaguchi, Yuichiro Kashiyama, Toshinobu Suzaki, Patrick J Keeling, Vladimir Hampl, Brian S Leander

**Author notes:** Corresponding authors: Anna Karnkowska, Yuichiro Kashiyama, Brian Leander. Three authors contributed equally to this work (Anna Karnkowska, Naoji Yubuki, Moe Maruyama). **Author Contributions:** A.K., N.Y., Y.K., P.J.K., V.H., and B.S.L. designed the research; N.Y., A.K., M.M., A.Y., and T.S. performed the research; N.Y., A.K., M.M., Y.K. and B.S.L. wrote the paper, with contributions from all authors.

## Abstract

Kleptoplasts are distinct among photosynthetic organelles in eukaryotes (i.e, plastids) because they are routinely sequestered from prey algal cells and function only temporarily in the new host cell. Therefore, the hosts of kleptoplasts benefit from photosynthesis without constitutive photoendosymbiosis. Here, we report that the euglenozoan *Rapaza viridis* has only kleptoplasts derived from a specific strain of green alga, *Tetraselmis* sp., but no canonical plastids like those found in its sister group, the Euglenophyceae. *R. viridis* showed a dynamic change in the accumulation of cytosolic polysaccharides in response to light– dark cycles, and ^13^C isotopic labeling of ambient bicarbonate demonstrated that these polysaccharides originate *in situ* via photosynthesis; these data indicate that the kleptoplasts of *R. viridis* are functionally active. We also identified 247 sequences encoding putative plastid-targeting proteins and 35 sequences of presumed kleptoplast transporters in the transcriptome of *R. viridis*. These genes originated in a wide range of algae other than *Tetraselmis* sp., the source of the kleptoplasts, suggesting a long history of repeated horizontal gene transfer events from different algal prey cells. Many of the kleptoplast proteins, as well as the protein-targeting system, in *R. viridis* were shared with members of the Euglenophyceae, providing evidence that the early stages in the endosymbiotic origin of euglenophyte plastids also involved kleptoplasty.

## Introduction

The endosymbiotic origin of mitochondria and plastids has been established with a wealth of evidence at different levels of biological organization. All of the plastids in eukaryotes, except those of the amoeboid *Paulinella chromatophora*, ultimately originated from a single endosymbiotic event with a cyanobacterium (1–4). However, the subsequent evolutionary history of plastids is much more complicated. Three lineages of photosynthetic eukaryotes, namely the Viridiplantae (green algae and land plants), the Rhodophyta (red algae) and the Glaucophyta, have retained the plastids acquired directly from the ancient cyanobacteria, called “primary endosymbiosis”. Several other eukaryotic lineages obtained plastids independently from one another by consuming either green algae or red algae, called “secondary endosymbiosis”. The eukaryotic groups with green secondary plastids are the Euglenophyta and the Chlorarachniophyta. Phototrophic cells in Stramenopiles, Alveolata, Haptophyta and Cryptophyta acquired their secondary plastid from red algal prey cells (3,5,6).

The number of different endosymbiotic events and the associated evolutionary processes that generated the diversity of plastids in eukaryotes remain contentious; however, the explanations for this history generally fall into one of two models that clarify the order of early stages that occurred before permanent plastids were established. The prevailing model infers a rapid integration step, in which a heterotrophic eukaryotic cell ingested an algal cell and maintained it as an endosymbiont rather than digesting it. The first step in this model is the development of a long-lasting predator-prey relationship between the host and its eventual photosynthetic endosymbiont. The establishment of mechanisms for metabolic exchanges must have then followed, which reduced the endosymbiont to an organelle through gene loss and gene transfer to the host nucleus (7,8). An alternative “targeting-ratchet” model for the endosymbiotic origin of plastids postulates that a protein-targeting system was established early to express proteins encoded in the host nuclear genome within the internalized plastid. This crucial evolutionary step drove the permanent integration of a plastid within its host cell (2). In this model, the gradual acquisition of critical genes by regular horizontal gene transfer events from prey cells, and the development of their targeting signals, was a prerequisite for the protein-targeting system that ultimately facilitated a permanent algal endosymbiont within the host cell (9).

Here, we investigated a potential model organism, *Rapaza viridis*, to better understand the evolutionary process of plastid acquisition. *Rapaza viridis* was first described in 2012 as the only mixotrophic euglenid known so far and is the closest sister taxon to the Euglenophyceae, a predominantly photoautotrophic group possessing a *Pyramimonas*-derived plastid (10). Behavioral data, ultrastructural data and molecular phylogenetic analyses of *R. viridis* demonstrated several intermediate morphological traits between those found in euglenophytes and phagotrophic euglenids (10). *Rapaza viridis* actively feeds only on a specific strain of *Tetraselmis* sp. using a reduced feeding apparatus and dies if deprived of either light or this strain of prey cells. In this study, we examined *R. viridis* and *Tetraselmis* sp. using transcriptomics, plastid genomics and time-course observations with transmission electron microscopy in order to evaluate the level of integration between the plastids and the host cell. We demonstrate that *R. viridis* has no canonical plastids, but is only transiently associated with kleptoplasts (11–18) (*SI Text*, Section 1.1), which are regularly acquired by consuming *Tetraselmis* sp. We also show that *R. viridis* expresses a series of nucleus-encoded proteins with apparent “plastid”-targeting signals that are homologous to those found in the Euglenophyceae and are probably transported into the kleptoplasts. Therefore, *R. viridis* provides a unique opportunity to examine the tenets of the targeting-ratchet model and the intermediate stages that occurred during the evolution of plastid-targeting systems in much greater detail than in any other previously reported organisms.

## Results and Discussion

### The first case of kleptoplasty in euglenozoans

The original description of *R. viridis* by Yamaguchi *et al*. (10) noted intermediate traits between those of phototrophic and heterotrophic (algivorous) euglenids and, in particular, a strong dependence on both modes of nutrition. It was inferred that *R. viridis* possesses canonical plastids and feeds actively only on specific prey, *Tetraselmis* sp., so the lack of either resource (light for photosynthesis or prey) was fatal (*SI Text*, Section 1.2). However, when we compared phagocytotic process of *R. viridis* with that of other algivorous cells, such as the phagotrophic euglenid *Peranema trichophorum*, we found that the cytoplasm of *R. viridis* lacked any typical digestive phagosome containing algal material. Among typical algivores, the light green color of the chlorophyll inside the phagosome fades to brown and the chlorophyll fluorescence disappears, suggesting the progression of digestion. However, this does not occur in *R. viridis*. This observation indicated that the prey, *Tetraselmis* sp., together with its plastids, is temporarily retained by *R. viridis* (10).

To verify this, we used the Sanger technique to sequence the 16S rRNA genes of the plastid genomes in a culture containing starved *R. viridis* (i.e., no trace of *Tetraselmis*) as well as from a unialgal culture of *Tetraselmis*. Unexpectedly, the single copies identified in each culture were identical and clearly matched the *Tetraselmis* sequence typical of Chlorodendrophyceae. Neither Euglenophyceae nor Pyramimonadales sequences were amplified from *R. viridis* (Fig. S1) (*SI Text*, Section 1.3). To confirm that the absence of Euglenophyceae 16S rRNA is not due to the low specificity of PCR, we assembled the whole plastid genomes from Illumina reads generated from the starved *R. viridis* culture and the prey *Tetraselmis* sp. culture. *R. viridis* data contained only one type of plastid genome which was identical to *Tetraselmis* sp. (Fig. S2; *SI Text*, Section 1.4). This result suggested that all of the cytoplasmic photosynthetic bodies of *R. viridis* were in fact kleptoplasts, or transient plastids snatched from its prey, *Tetraselmis* sp.

Our time-course observations of *R. viridis* using transmission electron microscopy revealed that the *Rapaza*-type plastid reported by Yamaguchi *et al*. (10) were derived from sequential transformation from the ingested *Tetraselmis*-type plastids (Fig. 1 and 2) (compare with *SI Text*, Section 1.2). Within 30 min after ingestion of a *Tetraselmis* sp. cell by phagocytosis of *R. viridis* (Fig. 1*A*–*C* and 2A; Movie S1), the plastid of *Tetraselmis* sp. was sorted and segregated from the other cellular material, which was eventually excreted from the cell (Fig. 1*D*–*F* and 2*B*). At this stage, the nucleus of *Tetraselmis* sp. Was eliminated (Fig. S3). In the following hours, the segregated kleptoplast was subdivided into smaller pieces (Fig. 1G–L and 2C), and constrictions formed an hourglass-like shape as observed with TEM (Fig. S4). Fission of the newly acquired kleptoplast was active 12–18 h after the ingestion event and resulted in the presence of ≤10 small pieces of kleptoplast in a single *R. viridis* cell. A large pyrenoid surrounded by polysaccharide grains was observed in the original plastid of *Tetraselmis*, but disappeared in an early stage before kleptoplast fission, while multiple smaller pyrenoids were formed that contain pyrenoid-penetrated thylakoids (Fig. 1*M*–*O*). At this stage, the kleptoplast displayed a three-membrane-bounded envelope, which is a typical feature of the *Rapaza*-type plastid (Fig. S5) (10). The early kleptoplast still contained the starch grains presumably formed inside the *Tetraselmis* plastid, indicating the *Tetraselmis* origin of the kleptoplast. However, in time, the starch grains gradually disappeared in the further subdivided pieces of the kleptoplast (Fig. 1*O*).

**Figure 1.**
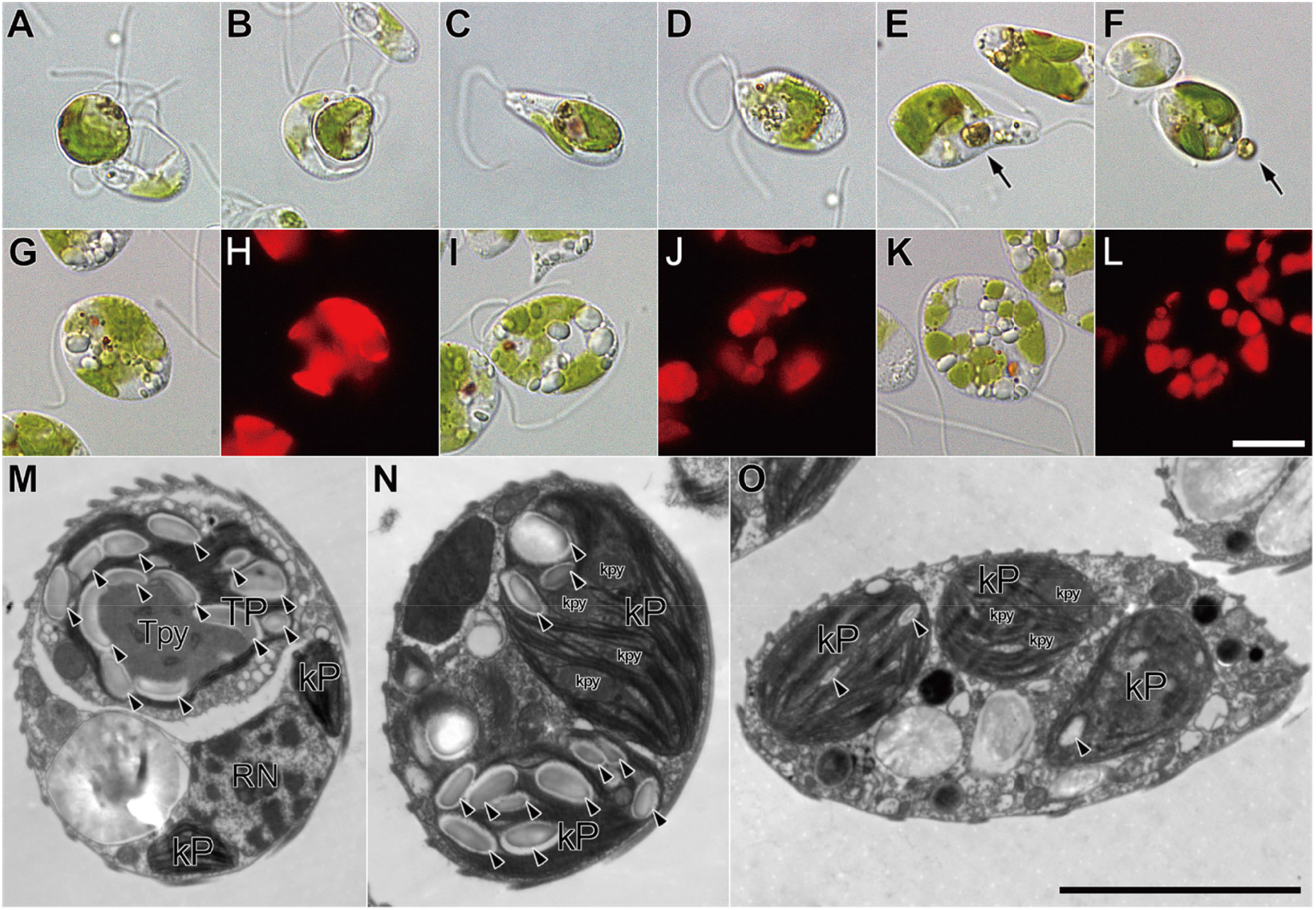
Time-series observations of *Rapaza viridis* using light and transmission electron microscopy (TEM) after the ingestion event. (*A*–*C*) *R. viridis* captures and engulfs cells of *Tetraselmis* sp., forming the phagosome. (*D*–*F*) Within 0.5–1 h of ingestion. *R. viridis* excretes the ingested cytoplasmic components of the *Tetraselmis*, except the plastid (arrow). The plastid is segregated and is the only component left in the cytoplasm of *R. viridis*. (*G*–*L*) The segregated plastid is subdivided into smaller pieces. Differential interference contrast images (*G*) 6 h, (*I*) 10 h, and (*K*) 14 h after the ingestion event. Corresponding fluorescent images show chlorophyll autofluorescence (excitation light: 400–440 nm) (*H, J*, and *L*). (*M*–*O*) Sequential TEM images of chemically fixed *R. viridis* cells (*M*) 0–1 h, (*N*) 6 h, and (*O*) 22 h after the ingestion event. (*M*) Immediately after the ingestion of a whole *Tetraselmis* cell containing its plastid (TP), which in turn contains a large pyrenoid (Tpy) and many polysaccharide grains (arrowhead). The cell of *Tetraselmis* is entirely contained within the phagosome of *R. viridis*. Nucleolus of *R. viridis* (RN) and old, previously captured kleptoplasts (kP) are visible. (*N* and *O*) The subdivided plastids gradually show the typical features of the *Rapaza*-type plastid (kleptoplast, kP), with diminishing intraplastidial starch grains (arrowhead) and the formation of multiple smaller pyrenoids that contain pyrenoid-penetrated thylakoids (kpy). White scale bar: 10 µm for (*A*–*L*); black scale bar: 5 µm for (*M*–*O*).

**Figure 2.**
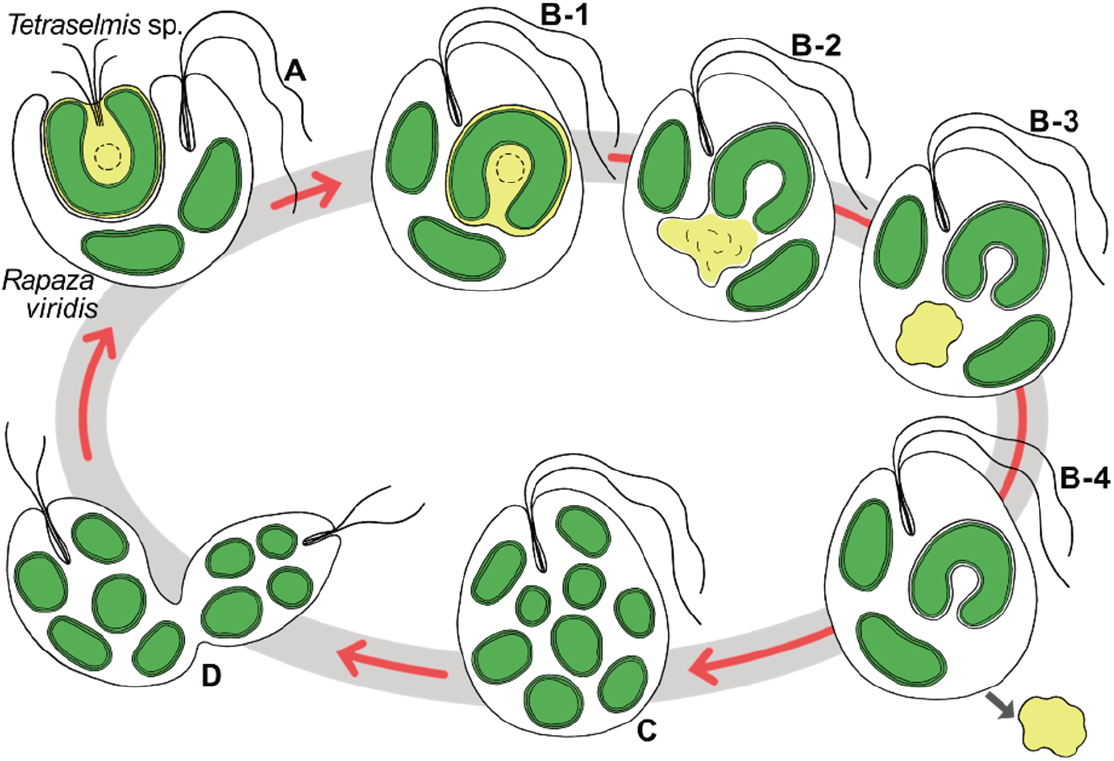
Schematics of the life cycle of *Rapaza viridis*. Because the kleptoplast is only temporarily functional, it must be replaced by the regular acquisition of fresh cells of *Tetraselmis* sp. (*A*) *R. viridis* ingests the entire cell of the *Tetraselmis* by phagocytosis. (*B*) *R. viridis* segregates the plastid of the ingested *Tetraselmis* from the other cytoplasmic components within the phagosome. The isolated plastid becomes tightly packed by the phagosomal membrane, which together with double membrane of the plastidial envelope, forms the triple membrane kleptoplastic envelope (*B-1* - *B-3*) (Fig. S4). The rest of the *Tetraselmis* sp. cytoplasm is quickly excreted from the cell (*B-4*). (*C*) The acquired kleptoplast is then subdivided within the cytoplasm of *R. viridis*. (*D*) Multiplied pieces of the kleptoplast are distributed to the daughter cells with no increase in the size of the kleptoplast beforehand or afterwards.

*Rapaza viridis* then initiated cell division in parallel with the later stages of kleptoplast fission (Fig. 2*D*), resulting in a threefold increase in cell number 11 days after the ingestion event (Fig. S6). Because the subdivided kleptoplasts were simply apportioned among the daughter cells of *R. viridis*, the number of the kleptoplasts per cell decreased as cell division was repeated. The growth continued until the total number of cells had increased 10-fold in 3 weeks, suggesting that 3–4 rounds of cell division occurred in *R. viridis* after ingesting a *Tetraselmis* cell. The total amount of chlorophyll per unit culture did not increase, despite the increased number of kleptoplast in cells, indicating that the amount of chlorophyll per cell of *R. viridis* decreased with increasing cell division (Fig. S6). This shows that the practical potential for photosynthesis of the kleptoplast in the cell of *R. viridis* did not increase after the ingestion event.

We then investigated whether kleptoplasts are actually used as the source of assimilates by *R. viridis*. We examined the daily dynamics of polysaccharide grains in the cytoplasm of *R. viridis* both microscopically and with chemical quantification. After kleptoplast acquisition was complete, polysaccharide grains accumulated in the cytoplasm of the daughter cells during the light period and reached their maximum just before the transition to the dark period. The quantity of accumulated polysaccharide grains decreased during the dark period and they were rarely observed just before the transition to the light period (Fig. 3 and Fig. S7). ^13^C isotope labeling experiments demonstrated that the polysaccharide grains in the cytoplasm were composed of photosynthetic products that were assumed to have been derived from the kleptoplasts. We added isotope-labeled bicarbonate (H^13^CO_3_−) to the medium during the light period and purified the polysaccharide grains from the cells just before the transition to the dark period for mass spectrometry analysis. Extensive incorporation of ^13^C into the polysaccharide grains was detected (Table S1). However, no accumulation of polysaccharide grains was observed in the cytoplasm of *R. viridis* when it was continuously shaded from light (Fig. 3*C*), suggesting that the polysaccharide granules did not derive from a heterotrophic carbon source but from the fixation of carbon dioxide.

**Figure 3.**
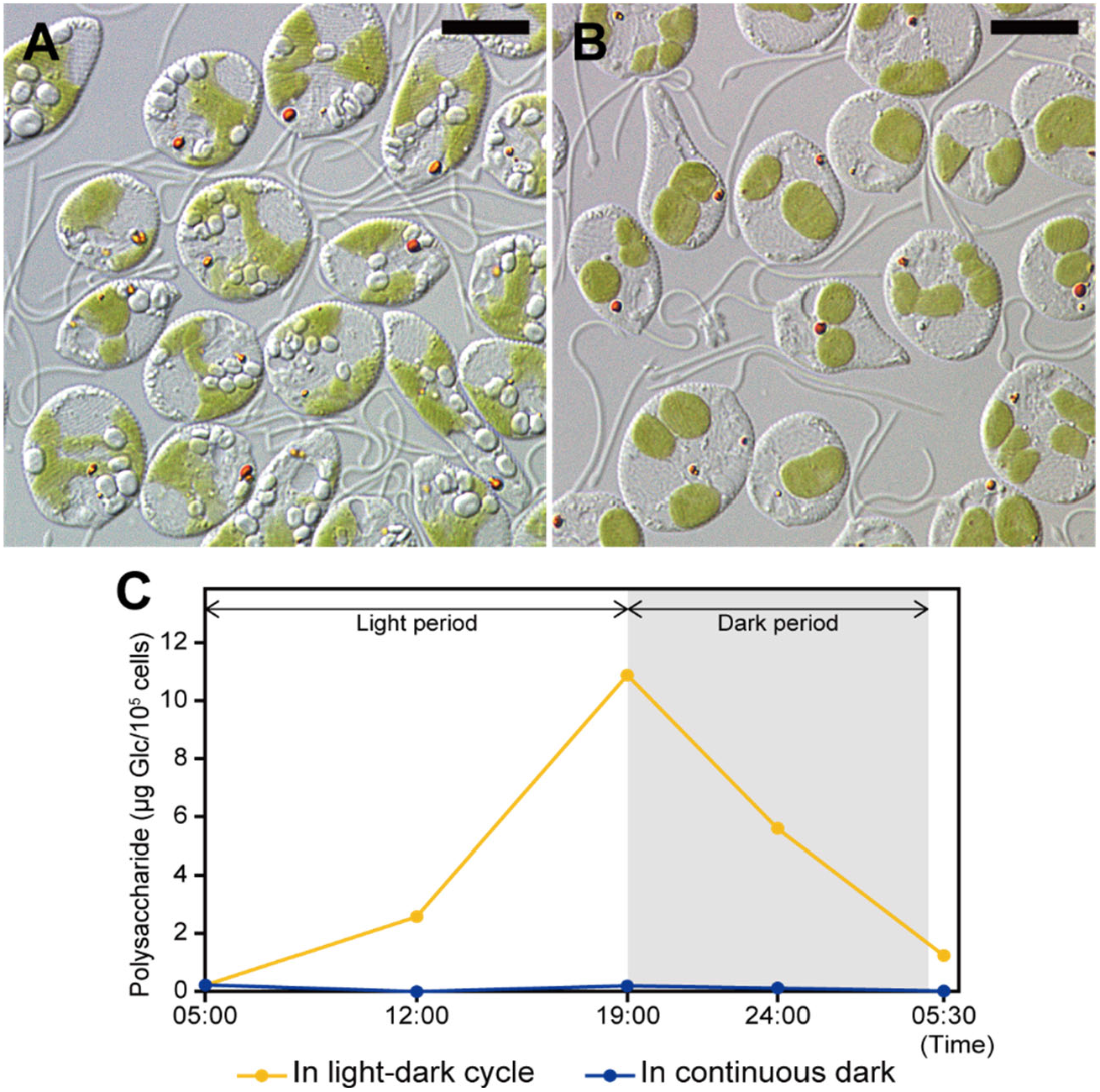
A diurnal cycle of the formation of the cytosolic polysaccharide grains in *Rapaza viridis*. (*A*) Polysaccharide grains resembling the paramylon granules of Euglenophyceae were visible and grew during the light period when *R. viridis* was cultured under a 14L:10D photoperiod. (*B*) However, the number of grains decreased during the dark period and had nearly disappeared by the end of the dark period. In contrast, no polysaccharide grains inside the kleptoplast were usually observed by TEM in fully established kleptoplasts, suggesting that polysaccharides only accumulated in the cytoplasm of *R. viridis*, unlike the accumulation of starch inside the plastid of *Tetraselmis* sp. (*C*) Quantitative data showed a correlation between the polysaccharide accumulation and the light intensity.

Neither genetic data nor morphological observations supported the presence of any *bona fide Pyramimonas*-type plastid in *R. viridis*. These data instead indicated the retention of plastids derived from its prey, *Tetraselmis* sp., providing the first evidence of kleptoplasty among euglenozoans. The increase in the number of kleptoplasts in the cells of *R. viridis* results from kleptoplast fission rather than the conventional division seen in other algae. Consequently, the multiplied kleptoplasts are distributed to the daughter cells with no increase in size. *R. viridis* appears to then become metabolically dependent upon the photosynthesis of these kleptoplasts, which must be supplemented by regular phagocytosis of fresh *Tetraselmis* prey cells.

### Evidence of nucleus-encoded genes for kleptoplasty

In the observations described above, the kleptoplasts in the cytoplasm of *R. viridis* remained functional after the elimination of the *Tetraselmis* nucleus in the earliest stage of kleptoplasty. This implies that at the molecular level, the host controls the kleptoplast functions to some extent with its own, nucleus-encoded proteins, to exploit the photosynthetic products of the kleptoplasts and, at least temporarily, maintain their function. To verify this possibility, we examined the transcriptome of *R. viridis* for genes encoding proteins that may be transported into the kleptoplasts.

We first searched for possible signal peptide sequence features similar to those known to be responsible for the translocation of nuclear-encoded proteins to *bona fide* plastids (19). As a result, we realized that the *R. viridis* sequence had features similar to the bipartite targeting system of Euglenophyceae, the closest relative of *R. viridis* (20, 21) (*SI Text*, Section 1.5). We predicted 2,241 putatively kleptoplast-targeted proteins based on the presence of the signal peptide (predicted with PrediSI or signalP) followed by the transit peptide (predicted with chloroP) and based on their homology to plastid-targeted proteins identified in *Euglena gracilis* (21). We then sifted through these candidate sequences using their predicted functions in plastids, based on their homology to either the predicted plastid proteome of *E. gracilis* (21) or the plastid-targeting proteins in the UniProt database (22). These methods identified 678 candidate genes (also see *SI Text*, Section 1.6). After the manual removal of potential contaminants, mitochondrial paralogues, and candidates with no transmembrane domain, 274 putative kleptoplast-targeted proteins were identified. We classified 166 of these as class I proteins with bipartite signal sequences and 73 as class II proteins with only a signal peptide (Table S2; *SI Text*, Section 1.5); we also annotated 35 proteins as transporters, constituting integral membrane proteins of the chloroplast envelope (Table S2). To validate the specificity of our approach on a negative control, we performed the same analysis on the predicted proteome of heterotrophic euglenid, *Rhabdomonas costata*, previously suggested to ancestrally lack chloroplasts (23). Here we predicted 2082 candidates with putative bipartite signals, which we then checked for contaminations, transmembrane domains and annotated their functions yielding 63 proteins (Table S3). Among those, 38 represent transporters (majority among them are ABC transporters and sugar transporters) with ambiguous localizations, and of the 26, only one (RCo055040, ornithine carbamoyltransferase, OTC) has a hit to a plastid-targeted protein in *Euglena gracilis*. Yet, the predicted N-termini signal in *R. costata* is much shorter than the one in the homologue from *R. viridis*, therefore we conclude this protein is a false positive hit.

If these proteins of *R. viridis* are truly targeted to the kleptoplast and, for classes I and II, function inside the *Tetraselmis*-derived kleptoplasts, we should identify a translocation complex in the triple-membrane envelope of the kleptoplast. Indeed, we identified one orthologue of each Tic21 (RV36988), Tic20 (RV1840) and Tic110 (RV32406) proteins in the transcriptome of *R. viridis*. Only two proteins (Tic 21 and Tic32 isoforms) have been identified in *E. gracilis* (24), and only Tic21 was confirmed in a proteomic study (21), suggesting highly reduced or derived translocation machinery in Euglenophyceae. Because Tic21 of *R. viridis* forms a well-supported clade with orthologues from Euglenophyceae (Fig. 4), it probably originated in their common ancestor, which likely possessed this unique translocation machinery and bipartite targeting signals on targeted proteins. However, other components (Tic20 and Tic110), which have together with Tic21 been proposed to form the inner envelope channel, are present specifically in *R. viridis* but not in Euglenophyceae. This suggests that these were either obtained later in the lineage of *R. viridis* or present in the common ancestor of *Rapaza* and Euglenophyceae and lost or diverged in Euglenophyceae. The independent origin of Tic20 is further supported by its relationship to ‘red-lineage’ (Fig. 4B) while the Tic100 is relative to ‘green-lineage’ (Fig. 4C). Overall, we conclude that *R. viridis* expresses nucleus-encoded proteins targeted to the kleptoplast and possess membrane translocons built from parts common with Euglenophyceae and other parts unique to *R. viridis*.

**Figure 4.**
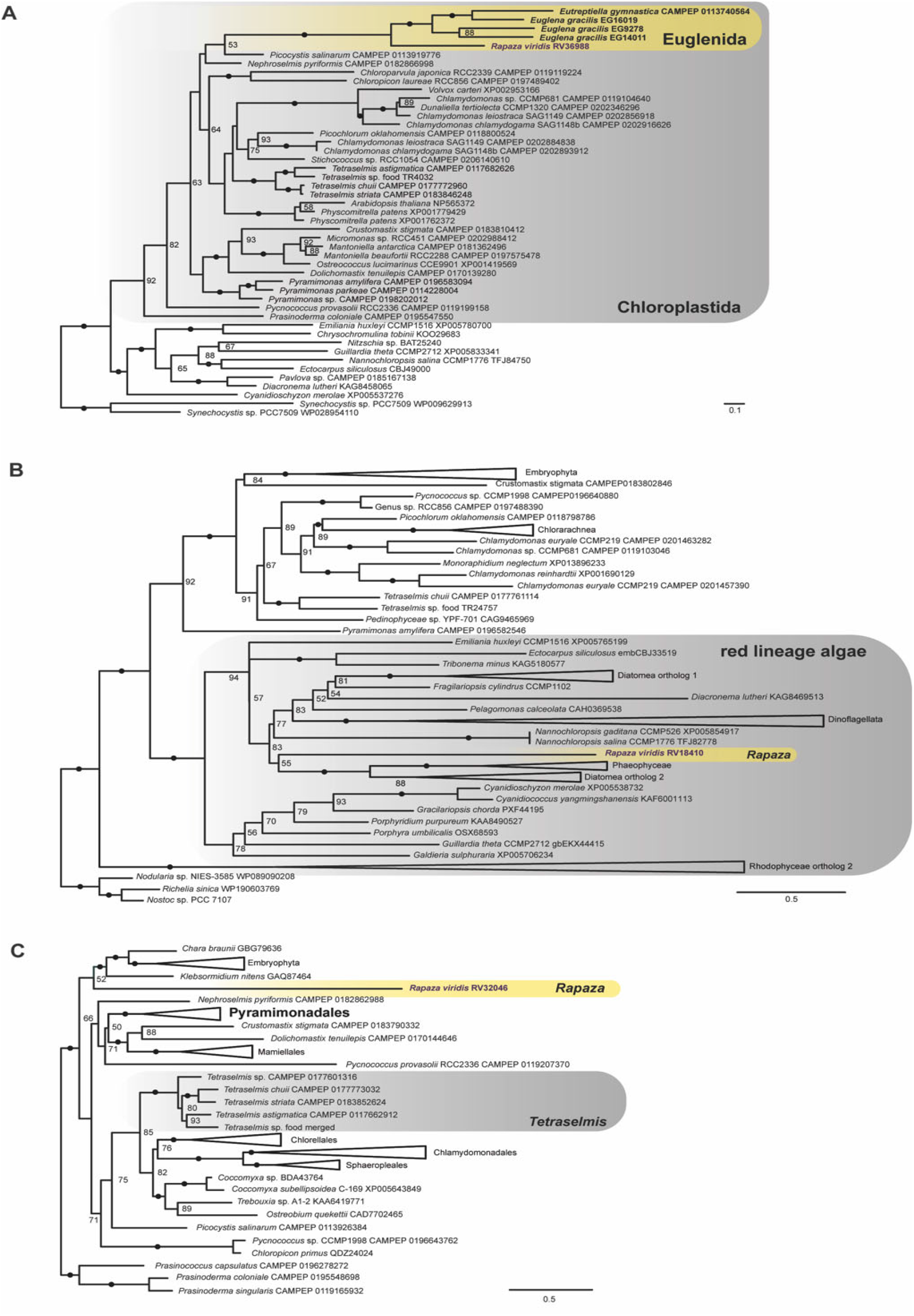
**A:** Phylogeny of Tic21 suggesting a shared ancestry of translocation machinery in *Rapaza viridis* and Euglenophyceae. **B:** Phylogeny of Tic20 suggesting a shared ancestry of this protein in *R. viridis* and ‘red lineages’. **C:** Phylogeny of Tic110 suggesting a shared ancestry of this protein in *R. viridis* and ‘green lineages’. Maximum likelihood tree of the Tic21 estimated with 1,000 rapid bootstrapping replicates in IQ-TREE. Photosynthetic euglenids are written in bold, and *R. viridis* is written in purple. Black circles (•) denote bootstrap support >95; support values below 50 are not shown.

### Diverse phylogenetic origins of kleptoplast-targeted proteins

To investigate the origins of proteins putatively targeted to kleptoplast, we calculated phylogenies for some of the candidate proteins and their orthologues from a custom protein sequence database. This database includes the genomes and transcriptomes of 77 algal lineages covering the major eukaryotic and cyanobacterial groups, in addition to representatives of Discoba. We also included the transcriptome of the prey *Tetraselmis* sp. to evaluate possible contamination.

Some of these genes in *R. viridis* encoding suspected kleptoplast-targeted proteins were confidently shown to originate from multiple green algal lineages other than *Pyramimonas* (Prasinophyta), from which the plastids of Euglenophyceae originated (25–27). Although many plastid-targeted genes grouped together within the *Tetraselmis* lineage, their sequences were never identical to those of the prey strain *Tetraselmis* sp. used in the culture system in this study. This observation strongly suggests a long-lasting, close interaction between *R. viridis* and *Tetraselmis*, which is inferred to have driven gene transfer from *Tetraselmis* to *R. viridis*. Clear such example is the chaperonin 60 beta subunit (Fig. 5*A*) in *R. viridis* clustered robustly with the *Tetraselmis* lineage of core chlorophytes, while homologues of Euglenophyceae formed a robust cluster with that of *Pyramimonas* spp., as anticipated. However, in other cases, including the ribulose-1,5-bisphosphate carboxylase/oxygenase activase gene, the sequence of *R. viridis* together with those of the Euglenophyceae formed a sister group (Fig. 5*B*). Other trees suggested a green algal origin of the plastid-targeting genes in all euglenophytes, but their exact origin was not possible to pinpoint, such as the gene encoding ftsH (Fig. S8*A*), a D1 quality control protease that is essential for photosystem II repair. Most interestingly, almost 30% of the genes encoding kleptoplast-targeted proteins were phylogenetically related to algal clades with red-algae-derived plastids, including Stramenopiles, Cryptophyta, and Haptophyta (i.e., collectively, the ‘red lineages’), rather than to any green algae. *R. viridis* homologues often clustered with Euglenophyceae (*Euglena* and *Eutreptiella*) (Fig. 6B and S8) in trees, suggesting gene transfer in a common ancestor of these euglenids (21, 28).

**Figure 5.**
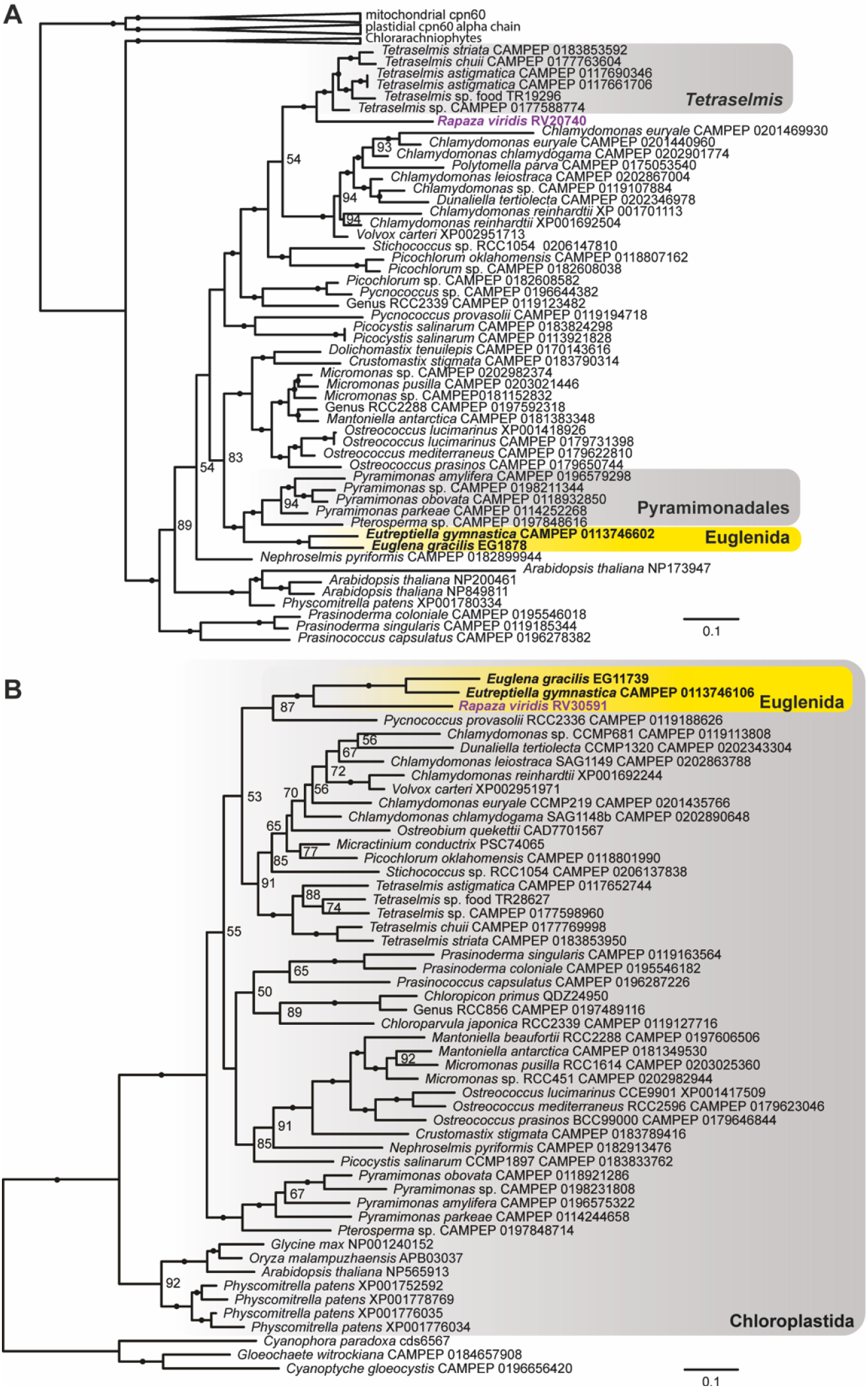
Diverse origins of the kleptoplast-targeted sequences in *Rapaza viridis*. Maximum likelihood trees were estimated with 1,000 rapid bootstrap replicates in IQ-TREE. Photosynthetic euglenids are highlighted in bold, and *R. viridis* is highlighted in purple. Black circles (.) denote bootstrap support >95; support values below 50 are not shown. **A:** Phylogeny of chaperonin 60 beta subunit (cpn60) showing sister relationship of *R. viridis* and *Tetraselmis* spp. **B**: Phylogeny of ribulose-1,5-bisphosphate carboxylase/oxygenase activase (RCA) showing sister relationship of clade of *R. viridis* and the Euglenophyceae.

**Figure 6.**
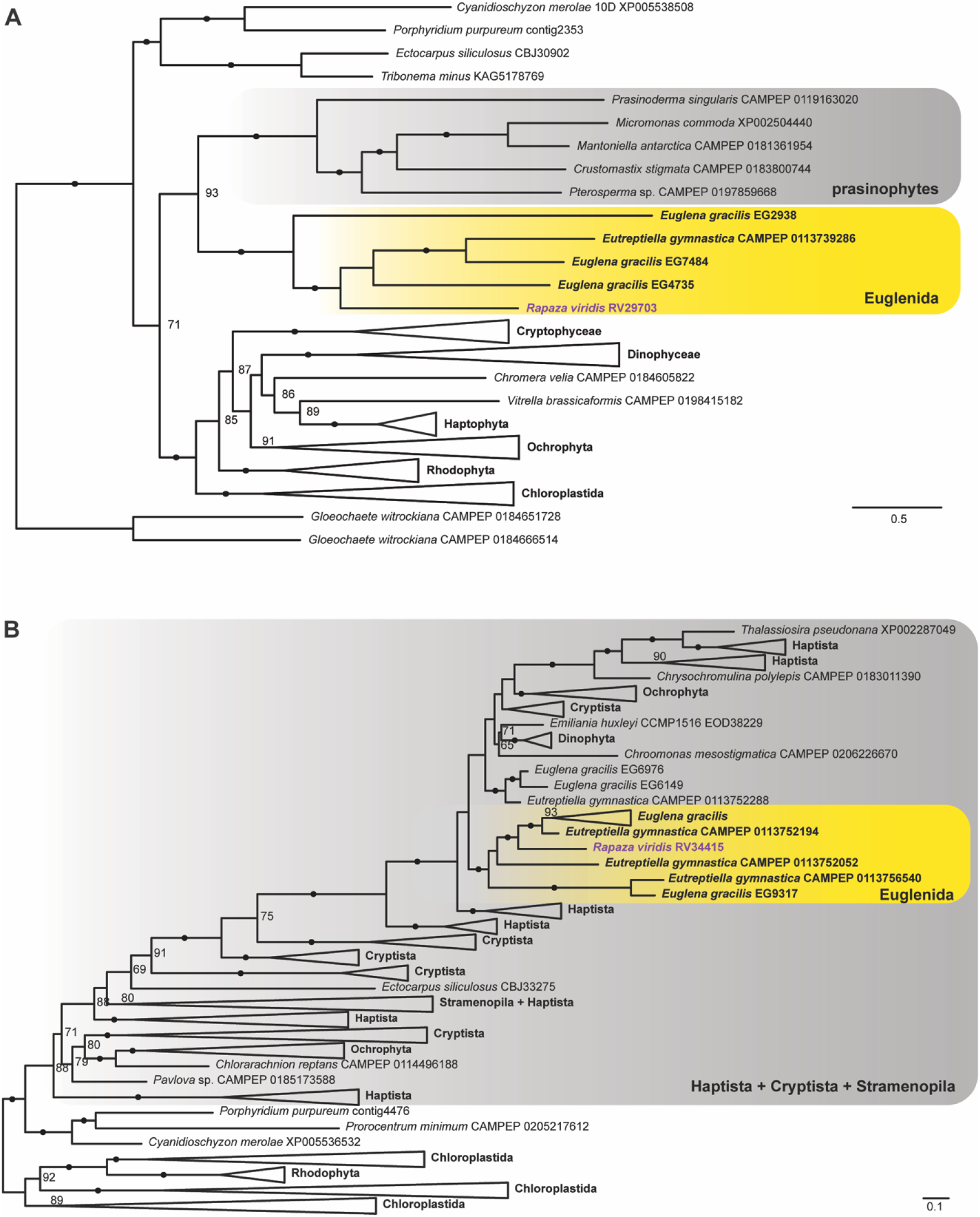
Maximum likelihood trees were estimated with 1,000 rapid bootstrap replicates in IQ-TREE. Photosynthetic euglenids are highlighted in bold, and *Rapaza viridis* is highlighted in purple. Black circles (•) denote boot strap support >95; support values below 50 are not shown. **A**: Phylogeny of plastidial glycolate translocator suggesting its green algal origin in *R. viridis* and Euglenophyceae. **B**: Phylogeny of triose phosphate/phosphate translocator (TPT) suggesting its transfer from the red algae-derived complex plastids to *R. viridis* and Euglenophyceae.

The observations described above support two inferences. First, these genes have been acquired by *R. viridis* in multiple horizontal gene transfer events involving a wide variety of algae, among which the core chlorophytes were predominant. This may reflect degrees of intimate interactions between *R. viridis* and these genetic sources, possibly as prey in kleptoplasty or algivory. Second, the shared presence of HGT genes is most likely attributable to their ancestral origin and must have been introduced into the lineage before the divergence of *R. viridis* and the Euglenophyceae. Notable such case is Tic21, a component of the TIC translocation machinery, which, together with the existence of similar bipartite targeting signals, indicates that the targeting system was established before the divergence of *Rapaza* and Euglenophyceae. Thus, it is also likely the pre-existing target system was involved in the establishment of the permanent plastid in Euglenophyceae. Our results agree with both the shopping bag hypothesis (29) and the red-carpet hypothesis (30), even though the origin of the plastids of Euglenophyceae is to be sought in the genus *Pyramimonas* (27).

Our data cannot fully resolve the relative timing of the symbiogenetic events, including the one that resulted in the *Pyramimonas*-derived plastid in Euglenophyceae. We infer that euglenids participated in more than a single symbiogenetic association at about the time when *Rapaza* and Euglenophyceae split. One scenario is that the last common ancestor of *R. viridis* and Euglenophyceae was a phagotrophic algivore without permanent plastids, which, however, might have already conducted kleptoplasty. Alternatively, one could assume the last common ancestor of *R. viridis* and Euglenophyceae had already established a permanent plastid derived from *Pyramimonas* together with a protein import system, which was subsequently eliminated in *Rapaza* that instead developed kleptoplasty with *Tetraselmis*. However, the former is a more plausible scenario considering (1) that *Pyramimonas*-derived genes are only minor in *R. viridis* and (2) that *Tetraselmis*-derived genes are rather abundant in Euglenophyceae; hence, the close association with *Pyramimonas* in the lineage of *Euglenophyceae* would have postdated the divergence. However, that scenario is less likely, due to (1) limited number of *Pyramimonas*-derived genes in *R. viridis* and (2) clear cases of *Tetraselmis*-derived genes among Euglenophyceae suggesting that the close association with *Pyramimonas* in the lineage of *Euglenophyceae* would have postdated the divergence.

### Metabolic transporters for exploiting the kleptoplast

We demonstrated that the products of photosynthesis from the kleptoplasts accumulate in the cytoplasm of *R. viridis*, which indicates the presence of a system for transporting organic molecules from the kleptoplast. We also observed that the starch grains inside the original *Tetraselmis* sp. plastid rapidly diminished in the earliest stage of kleptoplasty and did not form again in the later stages. *R. viridis* does not thrive in the absence of light, indicating its dependence on kleptoplasts for energy and the presence of transporters that deliver the necessary nutrients to the kleptoplasts. Therefore, we inferred that *R. viridis* expresses some metabolite transporters that pass through the envelope of the kleptoplast and transport the photosynthetic products. The importance of transporters for organellogenesis has been discussed previously in the endosymbiotic evolutionary model, where the targeting of transporters by the predator to its prey can facilitate the enslavement of the endosymbiont (2, 31).

We evaluated the metabolite transporters in the predicted proteome of *R. viridis*. Among the 274 predicted kleptoplast-targeting proteins, we identified 35 candidates that could function as metabolite transporters in the kleptoplast envelope (Table S2). The transporters seemed to be of exotic origin and transferred from prey cells representing either green algae (Fig. 6*A*; Table S2) or the red lineages (Fig. 6*B*). The phylogenetic trees indicated sister relationships between the homologues of *R. viridis* and Euglenophyceae, suggesting that the acquisition of the transporters predated their common ancestor.

### Targeting-ratchet model of kleptoplasty in *Rapaza viridis*

Kleptoplasty in *R. viridis* raises new insights into the order of events during symbiogenesis within the context of the targeting-ratchet model that differ from ideas proposed previously (32-34). Genes from both green algae and the red lineages have been reported in the genome of Euglenophyceae, which are expressed in the plastids and confer essential functions (21). Many of these genes are also shared with *R. viridis* (Figs 5 and 6), suggesting that these genes were already present in their last common ancestor. Furthermore, some of the shared genes include essential transporters for exploiting the plastids/kleptoplasts. Therefore, if the plastid in Euglenophyceae postdated the split from *R. viridis*, then the acquisition of their shared exotic genes and the plastid-targeting mechanism must have predated the first association of the common ancestor of Euglenophyceae with *Pyramimonas*. Alternatively, if a permanent plastid had been already established in their common ancestor, still the shared genes and the “kleptoplast”-targeting mechanism must have predated the modern association of *R. viridis* with *Tetraselmis*; little evidence of a close relationship between the common ancestor and *Tetraselmis* can be found in the repertoire of the shared genes. In either case, the acquisition of the gene and the establishment of the target mechanism had to precede the new symbiogenetic association.

This stands in contrast to the orthodox hypothesis for symbiogenesis inferred from previously reported intermediate organism (e.g., *Hatena arenicola*), in which endosymbiosis is inferred to have occurred before the transfer of endosymbiotic genes and the loss of the endosymbiont’s nucleus (34). Instead, plastid-targeting evolved early, probably during a long period of predation and kleptoplasty, and before the organelle become a permanent fixture of the cell. This has been also observed in a tertiary plastid origin (15), and these data suggest this may be a common route to endosymbiotic organelle origins. Additional comparisons of *R. viridis*, members of the Euglenophyceae, and potentially other yet to be discovered lineages that diverged near *R. viridis* will continue to clarify the order of evolutionary steps involved in the establishment of kleptoplasts and permanent plastids.

## Conclusions

Previous behavioral data, ultrastructural data and molecular phylogenetic analyses demonstrated several intermediate traits in *R. viridis* that fall between those in Euglenophyceae, and phagotrophic euglenids. Members of the Euglenophyceae possess secondary plastids originating from *Pyramimonas*-like green algae; however, many details about the origin and early evolution of euglenophyte plastids remain unclear. *R. viridis* is the only mixotrophic euglenid known and branches as the sister lineage to the Euglenophyceae, so *R. viridis* provides a unique opportunity to gather insights into the order of events associated with the origin(s) of plastids. We performed time-course TEM observations of kleptoplastid acquisition, biochemical analyses, plastid genomics and transcriptomics from *R. viridis* and its green algal prey *Tetraselmis* sp. Our data showed that *R. viridis* (1) represents the first case of kleptoplasty (i.e., functional plastids stolen from prey cells) in euglenozoans; (2) lacks the canonical plastids present in Euglenophyceae; (3) has nucleus-encoded genes for plastid-targeted proteins with euglenid-type targeting signals to the kleptoplasts; (4) has nucleus-encoded plastid-targeted proteins that originated from many different lineages of algae in addition to *Tetraselmis*; and (5) has plastid transporters to facilitate targeting of the proteins into the kleptoplasts. These data advance our understanding of the origin and evolution of plastids not only in the Euglenophyceae but also eukaryotes as a whole.

## Materials and Methods

### Cultures

The strains of *Rapaza viridis* (PRA-360) and *Tetraselmis* sp. (PRA-361) established by the authors of the original study (10) were also used in this study. These strains had been deposited in the American Type Culture Collection (Manassas, VA, USA) as PRA-360, but they subsequently died. They have now been deposited into the Microbial Culture Collection at the National Institute for Environmental Studies (Tsukuba, Japan) as NIES-4477 for *R. viridis* and NIES-4478 for *Tetraselmis* sp. Both strains were incubated in either f/2 medium (for genome and transcriptome sequencing) or Daigo IMK medium (Nihon Pharmaceutical Co., Ltd., Tokyo, Japan) for the remaining experiments, at 20 °C under illumination of 25–75 μmol-photons m^−2^ s^−1^ with a 14 h light/10 h dark photoperiod (14L:10D). The *R. viridis* culture was maintained by regularly inoculating fresh medium with it, together with three times the number of *Tetraselmis* sp. cells (1:3) every 2 weeks. After inoculation, the cells of *Tetraselmis* sp. were usually completely consumed by *R. viridis* within 12 h, resulting in a monospecific culture containing *R. viridis* exclusively. The monospecific culture of *Tetraselmis* sp. was maintained by inoculating fresh medium with an aliquot every 2–4 weeks.

To set the zero point for the time-course experiments involving serial light microscopy, TEM and chlorophyll analyses, fresh Daigo IMK medium was inoculated simultaneously with *R. viridis* cells and an equal number of *Tetraselmis* sp. cells (1:1) for rapid consumption, which was generally completed within 30 min. The cell numbers of both strains were counted with a particle counter/analyzer (CDA-1000, Sysmex, Kobe, Japan). For microscopic observations, the culture was incubated at 20 °C under 14L:10D (50 μmol-photons m^−2^ s^−1^), and the first dark period started 6 h after the inoculation. For the chlorophyll analysis, the culture was incubated at 20 °C under continuous light (50 μmol-photons m^-2^ s^-1^).

Quantification of polysaccharide accumulation was conducted with a culture of *R. viridis* 8–9 days after fresh Daigo IMK medium was inoculated simultaneously with *R. viridis* and three times the number of *Tetraselmis* sp. cells (1:3). The culture was then incubated at 20 °C under 14L:10D (50 μmol-photons m^−2^ s^−1^).

### Light microscopy

Differential interference contrast and bright-field images were generated with an inverted microscope (IX71, Olympus, Tokyo, Japan) equipped with a color CCD camera (FX630, Olympus) and the Flovel Image Filing System (Flovel, Chofu, Japan). Fluorescent images and the associated monochromatic bright-field images were generated with an inverted microscope (IX73, Olympus) equipped with a CMOS camera (ORCA-Flash 4.0, Hamamatsu Photonics, Hamamatsu, Japan) and cellSens Dimension v 1.18 software (Olympus). In general, the sampled cells were held between two glass coverslips (0.13–0.17 mm) for observation. To stain the nuclei of live cells, SYBR Green I (Lonza, Basel, Switzerland) was applied to the cultured cells before observations.

### Transmission electron microscopy

Chemical fixation was performed with a previously described method (35). Briefly, the cells were treated with glutaraldehyde and osmium tetroxide and then dehydrated through a graded series of ethanol before they were embedded in Spurr’s resin (Polysciences Inc., Warrington, PA). Rapid freezing and the freeze-substitution method were performed as previously described (36). The condensed cell cultures were cryofixed by impact-freezing onto a liquid-nitrogen-cooled copper block using a freezing device (VFZ-1, Japan Vacuum Device, Ltd., Mito, Japan) and were then freeze-substituted in acetone containing 1% OsO_4_ for 3 days. The specimens were returned to room temperature with a stepwise temperature rise before they were embedded in Spurr’s resin (Polysciences Inc.). Finally, ultrathin sections were stained with EM Stainer (Nisshin EM, Tokyo) and lead citrate before they were observed by TEM (Hitachi H7100, Hitachi Ltd., Tokyo, Japan).

### Quantitation of polysaccharide grains

The polysaccharide grains were extracted from *R. viridis* cells and purified as previously described (37). The precipitated insoluble polysaccharide was fully dissolved in 1 M NaOH and quantified with the phenol–sulfuric acid assay (38), with reference to standard glucose solutions.

### HPLC analysis

The pelleted cells frozen in liquid nitrogen were homogenized in acetone (1 mL) in an ice-cooled ultrasonication bath for pigment extraction. The acetone supernatant was immediately separated from the residue by centrifugation and directly injected into the HPLC apparatus for analysis. Analytical HPLC was performed with a liquid chromatograph system (Nexera X2, Shimadzu, Kyoto, Japan) coupled to a personal computer configured to run the Shimadzu LabSolution software (Shimadzu). Reversed-phase HPLC was performed according to the method of Kashiyama *et al*. (39).

### Isotope labeling experiment

In the labeled culture experiment, a 2 mM solution of isotopically labeled sodium bicarbonate (NaH^13^CO_3_; Cambridge Isotope Laboratories Inc., Tewksbury, MA) was added to a batch culture of *R. viridis* and to a culture blank medium (Daigo IMK) in a ratio of 1:1000. In the control culture experiment, a 2 mM solution of unlabeled NaHCO_3_ (Nacalai Tesque, Kyoto, Japan) was added to a batch culture of *R. viridis* in a ratio of 1:1000. An aliquot of the labeled blank medium was immediately dried *in vacuo*, and the carbon isotopic composition of the solid residue obtained was analyzed.

Subsamples (17 mL) of each of the labeled and control cultures were placed in a 50 mL vent-capped disposable culture flask and incubated at 20 °C for 9.5 h under continuous light (50 μmol-photons m^−2^ s^−1^). All the cells in each culture were then pelleted and frozen in liquid nitrogen. The purified polysaccharide grains were prepared from the frozen pellets using the method described above for polysaccharide quantification and their carbon isotopic compositions were analyzed by Shoko Science (Yokohama, Japan) and are expressed in the conventional δ-notation against Vienna Peedee Belemnite (δ^13^C (‰) ≡ 10^3^[(^13^C/^12^C)_sample_/(^13^C/^12^C)_standard_ − 1]).

### Amplification and sequencing of partial plastid 16S rDNA

The total DNA was extracted from cells harvested from 16-day-old cultures of *R. viridis* and *Tetraselmis* sp., with the Wizard Genomic DNA Purification Kit (Promega, Madison, WI), according to the procedure provided by the manufacturer. To enhance the extraction efficiency, the *Tetraselmis* sp. cells were bead-treated before the extraction procedure. The partial 16S rDNA in the plastid was amplified with PCR using a universal primer set that recognizes the known sequences of the family Chlorodendrophyceae, including *Tetraselmis*, and those of both Pyramimonadales and Euglenophyceae (TPE-16S_Fw: 5′-GTGCCAGCAGMYGCGGTAATAC-3′; and TPE-16S_Rv: 5′-TGTGACGGGCGGTGTGKRCAAR-3′). The amplified products were gel-purified with the Wizard SV Gel and PCR Clean-Up System (Promega) and then cloned into the pGEM-T Easy Vector (Promega). The inserted DNA fragments in the cloned plasmids were sequenced in both directions by Eurofins Genomics (Tokyo, Japan).

### Sample preparation for genomic and transcriptomic sequencing

The total genomic DNA of *R. viridis* and *Tetraselmis* sp. was extracted with the QIAmp DNA Micro Kit (Qiagen, Hilden, Germany). The total RNA was isolated from *R. viridis* and *Tetraselmis* sp. with the NucleoSpin RNA XS Kit (Macherey-Nagel, Düren, Germany). To avoid contamination with its food *Tetraselmis* sp., we extracted the DNA and RNA from 2-week-old cultures of *R. viridis*. We observed no *Tetraselmis* sp. cells in these cultures. We compared the small subunit ribosomal RNA (SSU rRNA) gene sequences with the genome database of *R. viridi*s and did not recover any *Tetraselmis* SSU rRNA sequences. This indicates that there was no contamination from *Tetraselmis* in the 2-week-old cultures of starved *R. viridis*.

Sequencing libraries were prepared and sequencing was performed on the Illumina MiSeq platform using 150-bp paired-end reads for the *R. viridis* genome, and with the Illumina HiSeq platform using 100-bp paired-end reads for the *R. viridis* at the University of British Columbia Sequencing and Bioinformatics Consortium, University of British Columbia, Vancouver, Canada. The *Tetraselmis* sp. genome and transcriptome was sequenced on the Illumina MiSeq platform using 200-bp paired-end reads at the BIOCEV, Charles University, Vestec, Czech Republic. The raw reads are available in the NCBI Short Read Archive (SRA) (for *R. viridis* Bioproject number: PRJNA901946, SRR22333116-SRR22333117 and for *Tetraselmis* sp. Bioproject number: PRJNA901983, SRR22356822-SRR22356821).

### Plastid genome and transcriptome assembly and annotation

Quality control of the Illumina MiSeq reads from the genomes of *R. viridis* and *Tetraselmis* sp. was performed with the FastQC tool v0.11.6 (40). Adapters, shortest reads (<36 bp), and poor-quality reads (mean Phred quality value of <15) were removed with the Trimmomatic tool v0.38 (41). The plastid genomes were pre-assembled with SPAdes v3.10.1 (42), and the plastid genes were identified in the assembled contigs using the BLASTX algorithm (43) and extracted. Contigs that contained the *rbc*L gene (encoding the large subunit of ribulose bisphosphate carboxylase/oxygenase) were extracted and used as the seeds for the final assembly of the plastid genomes with NOVOPlasty v2.6.3 (44). The chloroplast genome sequence have been deposited to the Dryad data depository (http://datadryad.org) accession doi:10.5061/dryad.37pvmcvpn. The plastid genome was annotated automatically with the cpGAVAS (45) and DOGMA tools (46), and manually corrected afterwards. Plastid genome map was created with the OGDraw tool (47).

Quality control of the Illumina HiSeq reads of the *R. viridis* and *Tetraselmis* sp. transcriptomes was performed with the FastQC tool v0.11.6 (40). Adapters, shortest reads (<36 bp), and poor-quality reads (mean Phred quality value of <15) were removed with the Trimmomatic tool v0.38 (41). The transcriptomes were assembled with Trinity v2.0.6 (48) and the proteins were predicted with TransDecoder v5.0.2 (https://github.com/TransDecoder/TransDecoder/releases/tag/TransDecoder-v5.0.2) and have been deposited to the Dryad data depository (http://datadryad.org) accession doi:10.5061/dryad.37pvmcvpn. The completeness of the transcriptomes was estimated with Benchmarking Universal Single-Copy Orthologs (BUSCO) v. 4.0.6 (49) with the eukaryote_odb10 dataset of 255 BUSCO groups. The predicted *R. viridis* proteome contained 87% complete BUSCOs and 3.5% fragmented ones, and 9.5% BUSCOs were missing. By comparison, the predicted proteome of *E. gracilis* (50) had 15.7% missing BUSCOs, suggesting that the *R. viridis* transcriptome was fairly complete. Similarly, the transcriptome of *Tetraselmis* sp. was estimated to be nearly complete, containing 83.9% complete BUSCOs and 7.8% fragmented BUSCOs, and 8.3% missing BUSCOs.

### Prediction of kleptoplast-targeting proteins in *R. viridis and R*.*costata*

The possible plastid-targeting proteins in the translated transcriptome (predicted proteome) of *R. viridis* and *R. costata* were predicted by searching for the typical bipartite plastid-targeting signal of euglenids (20) and by searching for orthologues of proteins identified in the plastid proteome of *E. gracilis* (21). The bipartite plastid-targeting sequences were identified as described by Novák Vanclová *et al*. (21) using a combination of SignalP v. (51) and PrediSI (http://www.predisi.de/) (52) to predict the signal peptides, and ChloroP v. 1.1 (53) to predict the chloroplast transit peptides after the *in silico* removal of the predicted signal peptides at their putative cleavage sites. The sequences were then truncated to a maximum length of 100 amino acid residues because the predictor searched for potential cleavage sites within the 100 most *N*-terminal residues. The preliminary dataset of *R. viridis* and *R. costata* plastid-targeting proteins (1563/1817 sequences) consisted of transcripts that had positive scores in SignalP + ChloroP, PrediSI + ChloroP, or SignalP + PrediSI + ChloroP analyses.

The translated transcriptome of *R. viridis* and *R. costata* was also compared with BLAST against the 1402 plastid-targeting proteins of *E. gracilis* (21), and the 874/276 candidate proteins with bidirectional best hits (e-value of 1e−20) were identified as plastid-targeting proteins. We combined both datasets, which resulted in 2241/2082 nonredundant candidate proteins, and annotated them automatically with BLAST at the *E. gracilis* plastid proteome (21) and UniProt databases (22). We manually annotated the proteins and excluded the redundant isoforms with the same annotation, mitochondrion-targeting proteins, and hypothetical and predicted proteins. We obtained a dataset of 678/261 annotated predicted plastid-targeting proteins and used it in the subsequent steps of the analysis.

### *N*-Terminal plastid-targeting sequences in *R. viridis and R. costata*

We evaluated the set of predicted *R. viridis* and *R. costata* chloroplast proteins (678/260 entries) for the characteristic *E. gracilis* plastid-targeting sequences. Potential membrane-spanning regions were identified with the hidden-Markov-model-based program TMHMM (54). The prediction was performed with sequences truncated to a maximum of 200 amino acids and that started with methionine, suggesting that they had complete *N*-termini. After the removal of too-short sequences, we obtained 274/64 sequences with at least one transmembrane domain (Table S2 and Table S3). Sequences with one transmembrane domain (73 proteins) were annotated as class II plastid-targeting proteins, and sequences with two transmembrane domains were annotated as class I (166 proteins) (Table S2). Sequences with three transmembrane domains (12 sequences) may represent those targeted across the thylakoid membrane, and were also classified as class I, according to Durnford and Gray (20). The remaining 35 proteins were annotated as transporters and constituted integral membrane proteins in the chloroplast envelope (Table S2).

### Determination of the evolutionary origins of plastid-targeting proteins

The sequences of *R. viridis* predicted to be plastid-targeting proteins with at least one transmembrane domain were used as queries for a BLASTP search against a custom database. The custom database consisted of predicted proteins, clustered at 90% similarity using Cd-hit (55), from the genomes and transcriptomes of 77 organisms, consisting of various groups of algae (including the transcriptome of *Tetraselmis* sp. sequenced in this study), several cyanobacteria, and species from the Discoba (including *E. gracilis* and *Eutreptiella* sp., representing the photosynthetic euglenids). The e-value cut-off used when collecting the sequences was 10^−3^. Each protein dataset was aligned using the MAFFT algorithm (with the default parameters) from the MAFFT package v7.271 (56). Regions of doubtful homology between sites were removed from the alignments with Block Mapping and Gathering with Entropy (BMGE) (57), with the default parameters. At this step, we discarded alignments with fewer than 70 sites after trimming (91 alignments) and those with fewer than 20 sequences (one alignment). The reduced protein datasets were realigned with the MAFFT-L-INS-I method in the MAFFT package, and then trimmed with BMGE (settings as previously described).

Maximum likelihood (ML) trees were constructed for the remaining 179 protein alignments using the IQ-TREE software v1.6.12 (58), with the evolution model automatically selected with the *-m TEST* parameter, and statistical support from 1000 rapid bootstrapping replicates (59). The trees were inspected manually to identify the relationships of the sequences from *R. viridis* with those from other euglenophytes and other algae included in the database, and to detect potential contamination and mitochondrial paralogues. To improve presented trees, we added additional homologs based on the NCBI database searches and manually inspected alignments to remove long branches and sequences shorter than half of the alignment and calculate the trees as described above.

## Acknowledgments

We thank Jun Kawahara, Taiki Koyama, and Tomomi Munekyo for their technical assistance; Dr. Masami Nakazawa (Osaka Prefecture University) for technical advice on experiments. This study was supported by grants to B.S. Leander from the Natural Sciences and Engineering Research Council of Canada (NSERC 2019-03986) and the Tula Foundation’s Hakai Institute; by a grant from the Gordon and Betty Moore foundation to P.J. Keeling (https://doi.org/10.37807GMBF9201), by the EMBO Installation Grant (grant number 4150) and the Ministry of Higher Education in Poland to A. Karnkowska; and by the Japan Society for the Promotion of Science (JSPS) KAKENHI to Y. Kashiyama (grant number JP18H03743, 21K19240).

## Notes

**Competing Interest Statement:** The authors declare no conflicts of interest.

### Competing Interest Statement

The authors have declared no competing interest.

