## Supporting Information for "Euglenozoan kleptoplasty illuminates the early evolution of photoendosymbiosis"

**Supporting Information for**
### Euglenozoan kleptoplasty illuminates the early evolution of photoendosymbiosis

Anna Karnkowska, Naoji Yubuki, Moe Maruyama, Aika Yamaguchi, Yuichiro Kashiyama, Toshinobu Suzaki, Patrick J Keeling, Vladimir Hampl, and Brian S Leander

*Corresponding authors: Anna Karnkowska, Yuichiro Kashiyama, Brian S Leander


**This PDF file includes:**

Supporting text
Figures S1 to S8
Table S1
Legends for Movies S1
SI References

**Other supporting materials for this manuscript include the following:**

Tables S2-S3
Movies S1

**Supporting Information Text**

**1. Summary of previous works on kleptoplasty.** During the course of the plastid acquisition through algal endosymbiosis, the existence of the intermediate states before establishment of permanent plastids have been speculated. The kleptoplasty is one of the forms of acquired phototrophy with non-canonical plastids and are studied in anticipation that those might help to reconstruct early stages in the evolution of plastids (1, 2). The hosts retain the kleptoplasts (stolen temporary chloroplasts) within their non-photosynthetic cell/tissue but only temporarily. In some case such as of dinoflagellates, kleptoplasts are co-retained with the algal nuclei (3, 4); however, in other cases, kleptoplasts are obviously uncoupled from the control by the algal nuclei that are removed upon acquisition of the kleptoplasts. Thus, lacking inherent ability to fully synthesize, repair, and control the photosynthetic machinery, the host needs to replace the temporary kleptoplasts from engulfed algal preys in each generation and/or repeatedly in the same generation.

Beside the herein reported Euglenozoan case, various degrees of acquired phototrophy as kleptoplasty have been observed in some distinct lineages in the tree of eukaryotes (5) including various dinoflagellates (3, 4, 6-9), ciliates (10), foraminiferans (11), a katablepharid (12), as well as metazoans (sea slugs) (13-19). Amongst them, well-studied kleptoplasty is known from a sea slug *Elysia chlorotica* that maintains plastids from a stramenopile alga *Vaucheria litrea* for up to 10 months (13). Although many early studies reported kleptoplast targeting sequences in the *Elysia* nucleus (14-16), the gene transfer from the alga for the kleptoplasts to the host *Elysia* nucleus genome as well as the protein targeting from the *Elysia* nucleus to the kleptoplasts remain contentious issue. Only trace levels of the putative plastid-targeted genes detected led to conclude that the microalgae RNA was contaminated (17). In addition, there was no plastid gene that could be recognized from the transcriptome data of *Elysia* eggs (18). Finally, recent transcriptomic and genomic studies demonstrated no positive evidence for LGT of plastid genes in the sea slugs (19).

Dinoflagellates offers several examples of kleptoplasty. A dinoflagellate, *Dinophysis acuminata*, has kleptoplasts with a complicated origin. Primarily, a ciliate, *Mirionecta rubrum*, feeds on a cryptophyte algae, *Germinigera cryophile*, and transiently acquires their plastids in the cell as kleptoplasts. Then, the dinoflagellate, *D. acuminata*, sequesters the cryptophyte plastids by feeding on *M. rubrum*. Wisecaver and Hackett (6) identified five putative kleptoplast-targeting sequences encoded in the nuclear genome in *D. acuminate*, where only PsbM (photosystem II subunit M) was presumably originated from the cryptophyte algae in their phylogenetic analyses. Since *Dinophysis* is thought to have lost its plastid during the evolution of this lineage, the origin of the other four genes was suggested to be its photosynthetic ancestor. Thus, the original PsbM of the photosynthetic ancestor might have been replaced with the one drived from the cryptophyte.

One of the well described examples is an obligatory kleptoplastic Ross Sea dinoflagellate (RSD) belonging to Kareniaceae. Two genera of Kareniaceae harboring stable tertiary plastids derived from haptophytes (7), and kleptoplastidic RSD also obtains kleptoplasts from its haptophyte prey, *Phaeocystis antarctica* (8). The analysis of closely related species with varying degrees of plastid integration not only proved the existence of kleptoplast-targeted proteins, but also indicated that the RSD and its closely related photosynthetic ancestors likely had already acquired some genes for plastid/kleptoplast-targeted proteins (9). This in turn suggested that the establishment of a protein-targeting system should have precede full integration of the novel plastid.

**2. The original description of *Rapaza*.** Based solely on the morphological observations, Yamaguchi *et al.* (20) described presence of original plastids that are to be distinct from that of the prey *Tetraselmis*, because (1) their ultrastructure of the “plastids” resembled those of Euglenophyceae more than that of *Tetraselmis*, (2) the “plastid” envelope of *Rapaza* contained three membranes as seen in Euglenophyceae, instead of two membranes in *Tetraselmis* and other green algae, and (3) polysaccharide grains were stored outside the “plastids” in *Rapaza* as in Euglenophyceae, but inside the plastid in *Tetraselmis* as in the other green algae.

**3. Phylogenetic relationships of *Rapaza viridis* to Euglenophyceae and origin of the euglenophycean plastid.** Origin of the secondary plastid of Euglenophyceae (i.e., phototrophic euglenids) has been studied through the combination of molecular phylogenetics, genomics and morphological investigations, which offered a well-supported and consistent view of the *Pyramimonas* plastid as the closest extant relative to the euglenophyte plastid (21, 22). Both morphological and molecular phylogenetic data have shown beyond any serious doubt that the Euglenophyceae comprises a monophyletic subgroup nested within a major non-photosynthetic trunk of Eugleinda (20, 23). Therefore, the phylogenetic null hypothesis for the origin of *Rapaza*'s canonical plastids, if presented any as described in the original description by Yamaguchi *et al*. (20), must also be *Pyramimonas*. Although both *Tetraselmis* and *Pyramimonas* are green algae and have similar morphology in light microscopy, they are phylogenetically very distant: *Tetraselmis* sp. belongs to Chlorodendrophyceae in the core chlorophytes, whereas *Pyramimonas* is related to Pyramimonadales that is a rather basal lineage of Chlorophyta (24, 25)

**4. Plastid genomes from *Rapaza* and *Tetraselmis*.** The size of circular plastid genome identified in *Rapaza viridis* at 98,479 bp is comparable with green algae, such as *Pyramimonas parkeae* at 101,605 bp (26) and *Tetraselmis* sp. CCMP881 at 100,264 bp (27). Those plastid genomes contain large and small single copy regions (LSC and SSC) and two inverted repeat regions (Fig. S2), much like plastid genomes of Chlorophyta and several lineages of Euglenophyceae (28). The plastid genome in *R. viridis* contains 70 protein coding genes, 26 tRNA genes, three rRNA genes and eight hypothetical conserved ORFs. Two genes (psaA and psbC) contain one intron each.

**5. General features of signal peptide in Euglenophyceae.** The positions of the signal peptides of Euglenophyceae are estimated between residues 9-60 (29); the signal peptide is followed by a plant-like plastid-targeting signal (transit peptide) for a possible TOC/TIC-like translocon in the inner membranes of the plastid envelope. A recent bioinformatic and proteomic analyses of euglenophytes confirmed plastid localization of only three isoforms of Tic21, while other components of the complex as well as all TOC subunits are missing or are highly divergent (30, 31). In some plastid-targeted proteins, a characteristic transmembrane domain of the stop transfer signal (STS) has been detected between residues 97-130. Based on the presence or absence of transmembrane domains, three types of plastid-targeted proteins have been described in the euglenophyte: Class I with two hydrophobic domains (SP and STS), Class II with only one transmembrane domain (SP) and the third type without hydrophobic domains found in some plastid-targeted proteins.

**6. Possible false negatives in the sifting of the candidate sequences.** In almost half of the primary candidates, any transmembrane domain was failed to be identified. However, we suspected that these might be false negatives: incomplete sequences that do not exhibit the typical plastid-targeting signal. For example, such a fraction of the false negatives was estimated to constitute up to 16% of the plastid proteome of *E. gracilis*. Thus, we were not able to classify those into three categories and therefore excluded those sequences form the detailed analyses of the kleptoplast-targeting signal and phylogenetic analyses. As a result, contrary to the general principle describing kleptoplasts as temporal plastids stolen from the food algae with no control by the host nucleus, we identified 274 genes in *Rapaza* inferred to encode proteins targeted to the kleptoplast.

**Fig. S1.** Alignment of the partial 16SrRNA genes in the plastid genomes sequenced from *Rapaza viridis* and *Tetraselmis* sp. (the prey of *R. viridis*), and representative sequences obtained from the GenBank database for Euglenophecеae (*E. gymnastica* NIES-381 [GenBank: FJ719672.1]) and Pyramimonadales (*Pyramimonas parkeae* CCMP 726 [GenBank: FN563104.1]) as references. The shaded area in the alignment indicates locations that are identical to the *R. viridis* sequence. The sequences from *R. viridis* and the prey *Tetraselmis* were exactly identical, and clearly different from those of Euglenophyceae and Pyramimonadales, indicating the absence of a Euglenophyceae-type plastid genome in *R. viridis*. Therefore, all the “plastids” present in the cell of *R. viridis* are inferred to be exclusively *Tetraselmis*-derived kleptoplasts.

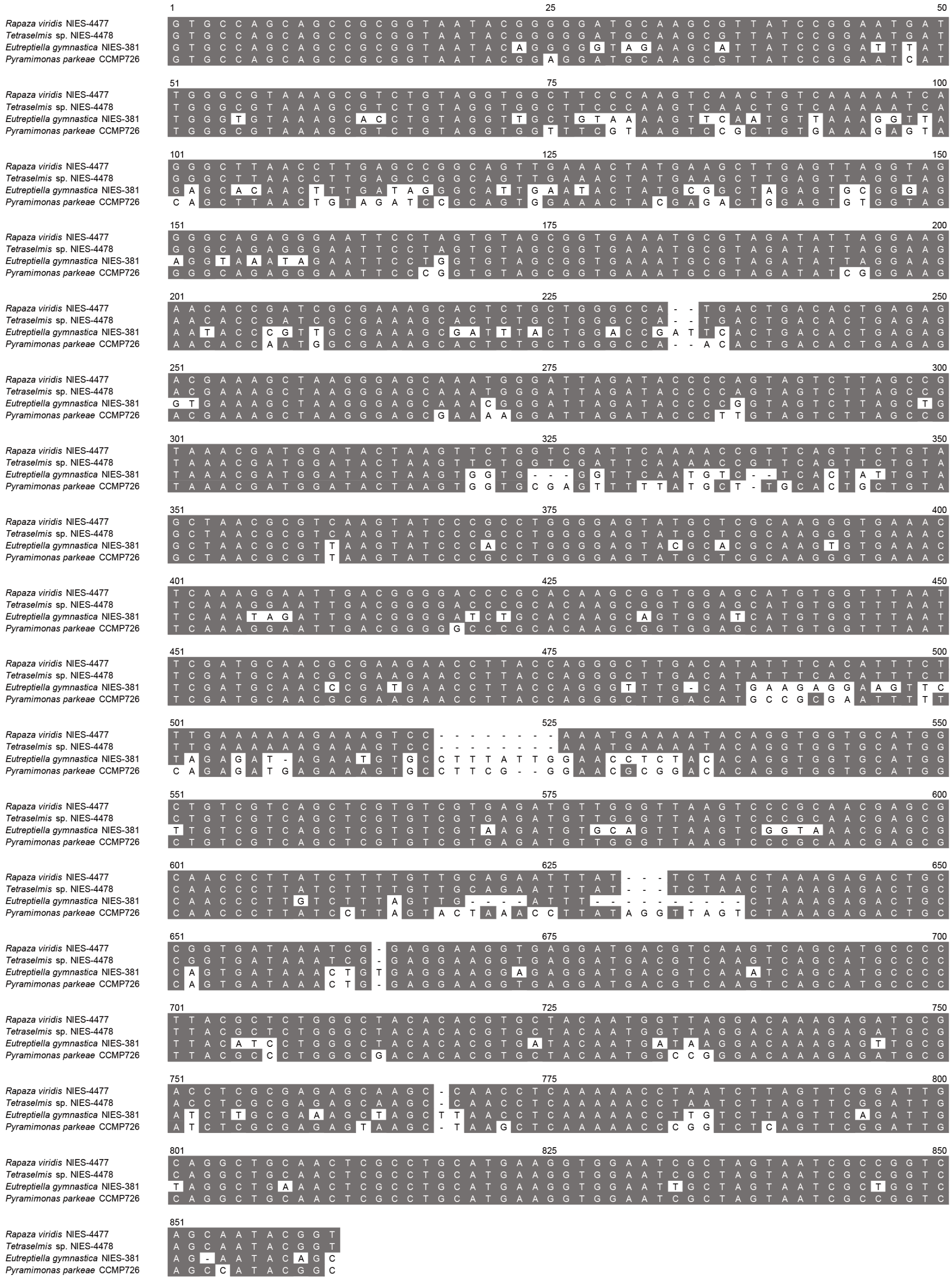

**Fig. S2.** Chloroplast genome map of *Tetraselmis* sp. obtained from the *Rapaza viridis* culture, showing the large single copy (LSC) and small single copy (SSC) regions and two inverted repeat regions (IRA and IRB, respectively).

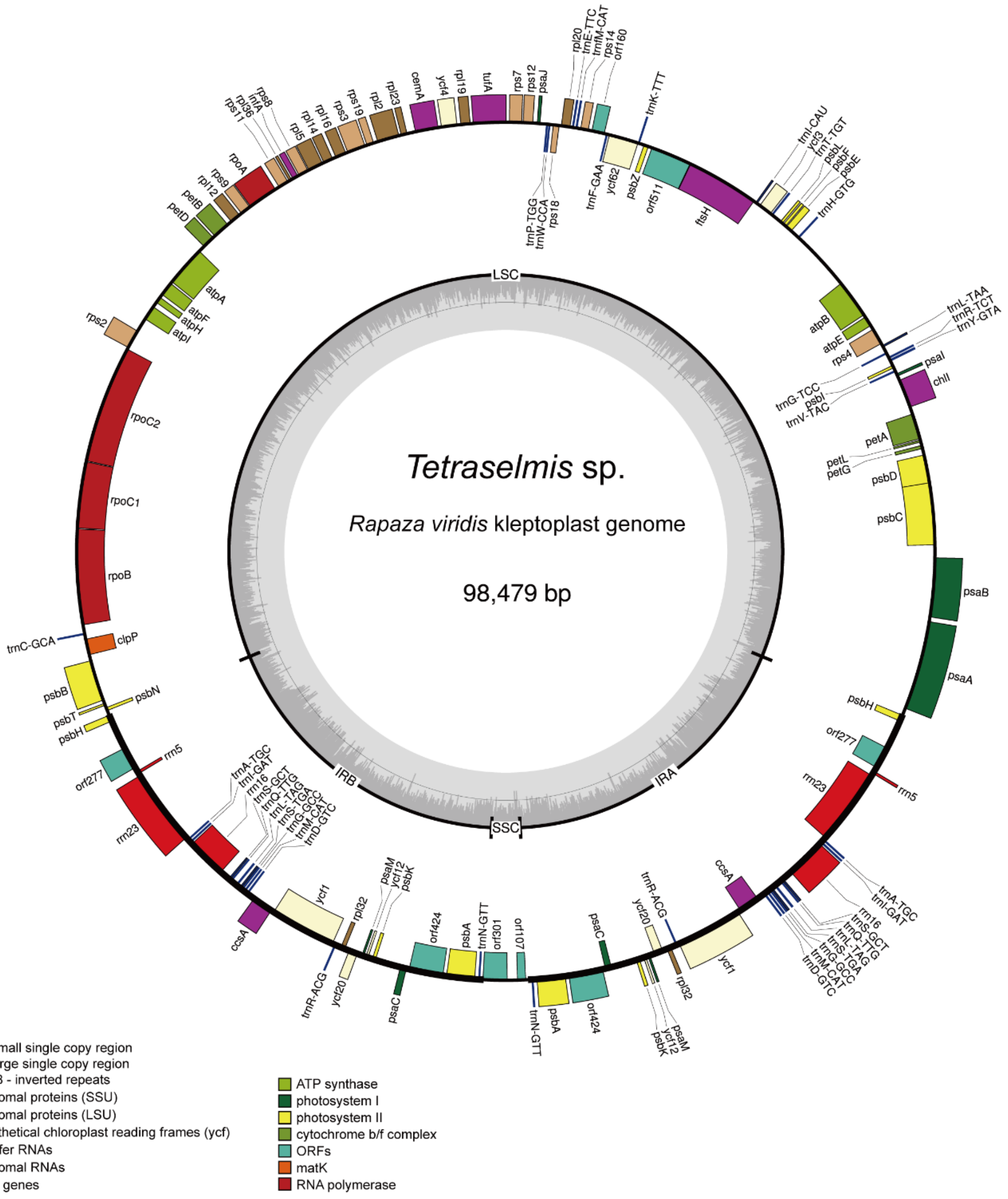

**Fig. S3.** Nuclear staining images of live *Rapaza viridis* 2 h after the ingestion event. (*A*) A bright-field image and (*B*) a corresponding multichromatic fluorescent image. Green fluorescence from SYBR-Green-stained nucleic acid indicates the presence of only the *R. viridis* nucleus, with no nucleus from the ingested *Tetraselmis* sp., suggesting that the nucleus of *Tetraselmis* is eliminated soon after ingestion. Red fluorescence is chlorophyll autofluorescence from the plastids (kleptoplasts). Scale bar: 10 μm.

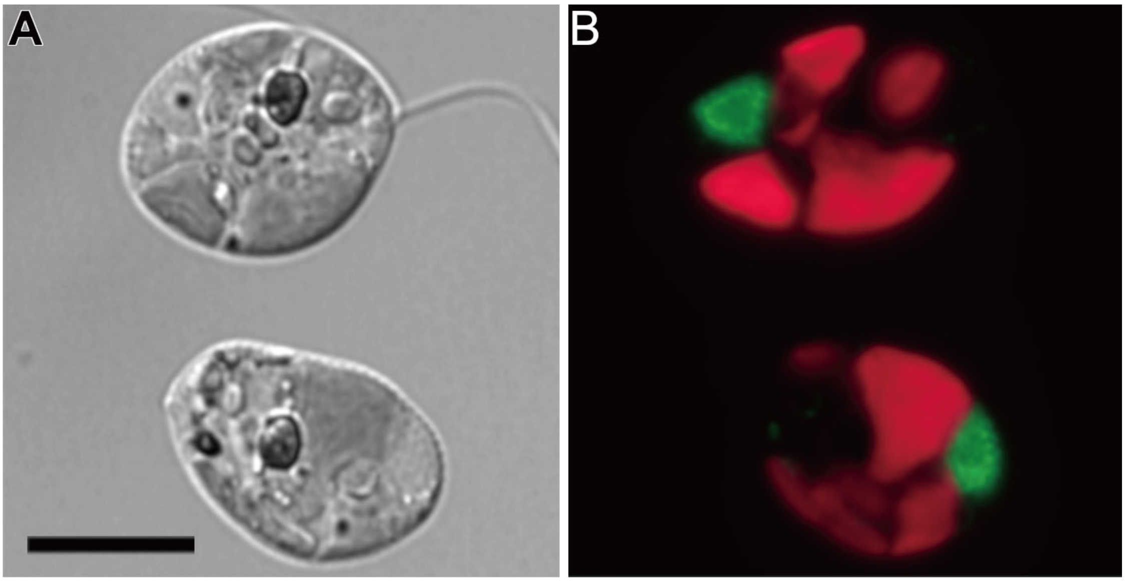

**Fig. S4.** TEM images of a *Rapaza viridis* cell displaying plastid fission. (*A*) Constricting structures creating an hourglass-like form were observed in cells fixed with the quick-freezing and freeze-substitution method. (*B*) Inside the constriction was typically densely stained (arrow). The constricting structure was not maintained after conventional chemical fixation (data not shown). Scale bars: 1 μm. kP: kleptoplast; Rc: *Rapaza viridis* cytosol.

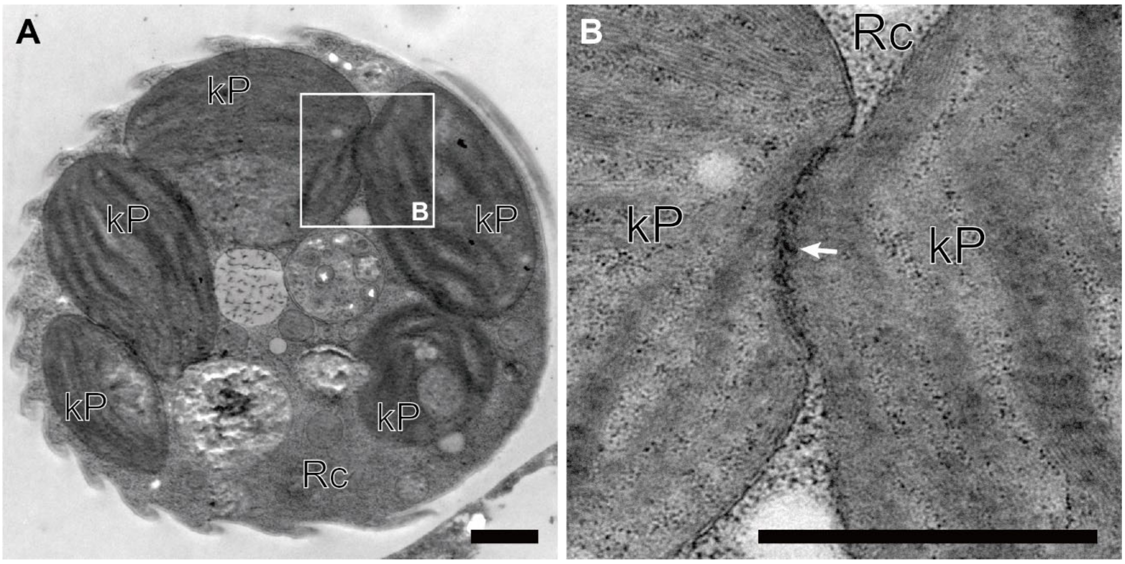

**Fig. S5.** TEM observations of the envelope bounding the subdivided plastids in an early stage of fission. (*A*) Presence of three membranes, as in Euglenophyceae, was typically detected in samples prepared with the chemical fixation method. We infer that the inner two membranes (arrow) correspond to the double membrane of the plastid of *Tetraselmis* sp., and the outermost membrane (arrowhead) that originated in the phagosomal membrane of *Rapaza viridis*. (*B*) In samples prepared with the quick-freezing and freeze-substitution method, the envelope was tightly stacked (arrow), indicating active elimination of the redundant intermembrane space. Scale bars: 200 nm. kP: kleptoplast; Rc: *Rapaza viridis* cytosol.

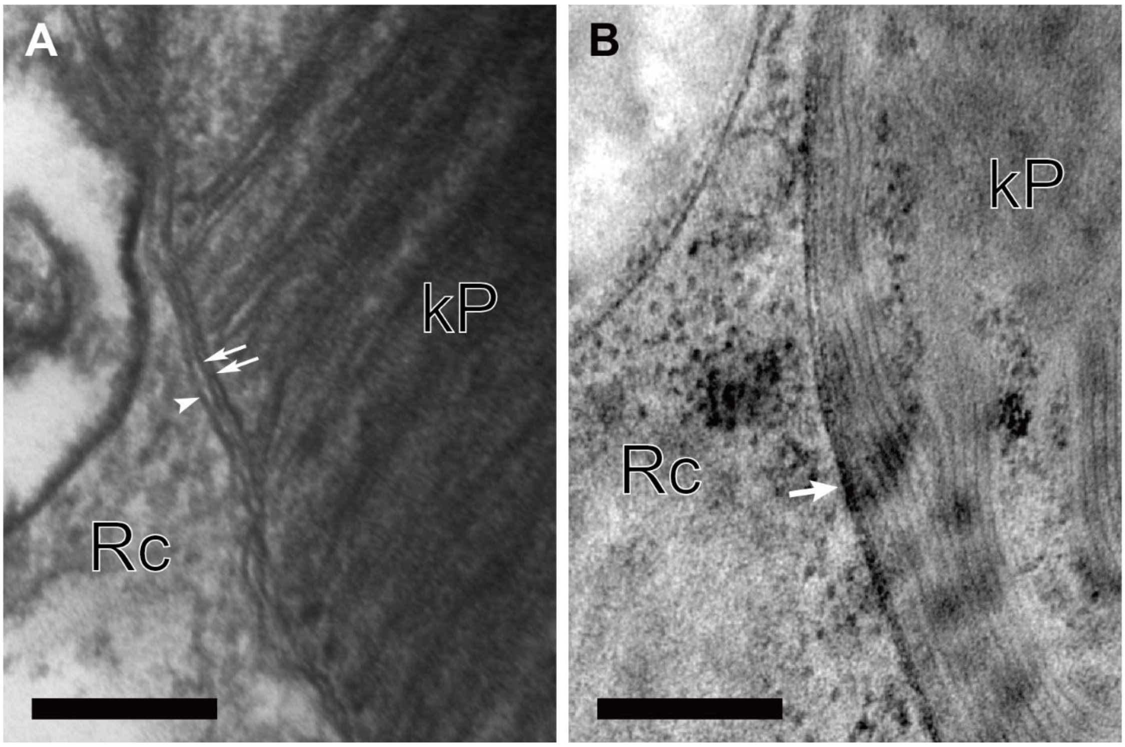

**Fig. S6.** Comparisons of changes in cell density during batch culture and the chlorophyll contents after the simultaneous ingestion of *Tetraselmis* sp. by *Rapaza viridis* (*n* = 3). (*A*) Changes in *R. viridis* cell density and chlorophyll *a*/*b* concentrations per unit culture volume. (*B*) Changes in *R. viridis* cell density and chlorophyll *a*/*b* content per cell. Cell density gradually increased after the ingestion event, whereas the chlorophyll content in the culture did not increase in parallel. Therefore, the calculated chlorophyll content per *R. viridis* cell decreased as cell division progressed, indicating that no new chlorophyll was produced in the kleptoplast.

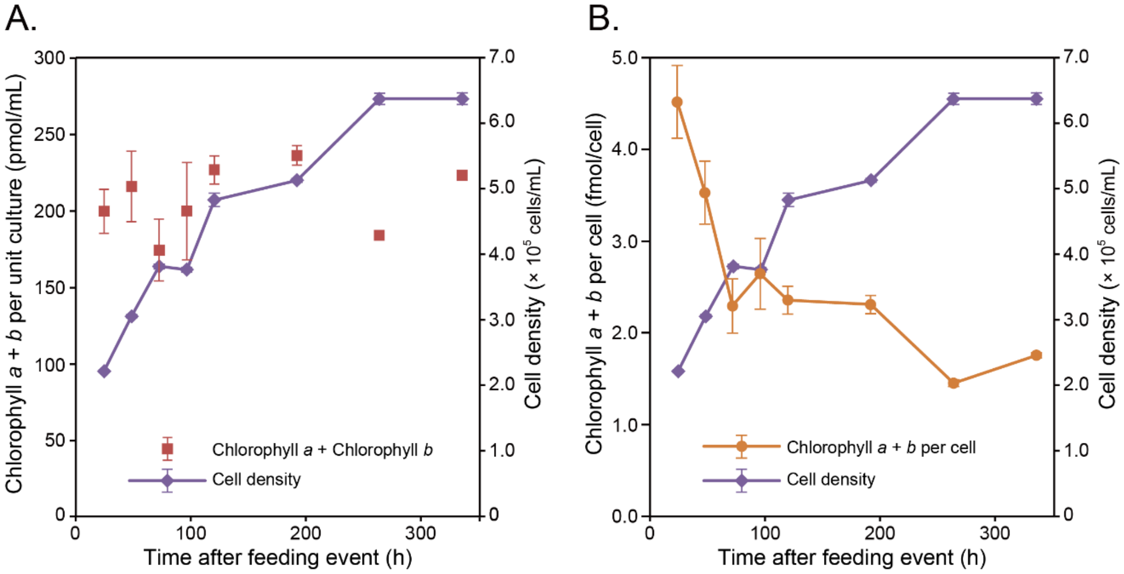

**Fig. S7.** Time-series of microscope images of the growth and decay of the cytosolic polysaccharide grains in *Rapaza viridis* in the 24-h experiment shown in Fig. 3C. (*A*) Cells immediately before onset of the light period at 05:00, in which few polysaccharide grains were observed. (*B*) Series of cells incubated in the light for 16 h (05:00–21:00), and then in the dark for 8 h (21:00–05:00). (*C*) Series of cells incubated in continuous dark for 24 h. Because the cytosolic polysaccharide grains only grew in the light, we infer that *R. viridis* accumulates the photosynthetic products as polysaccharide grains in the cytoplasm during the day and consumes them by respiration at night.

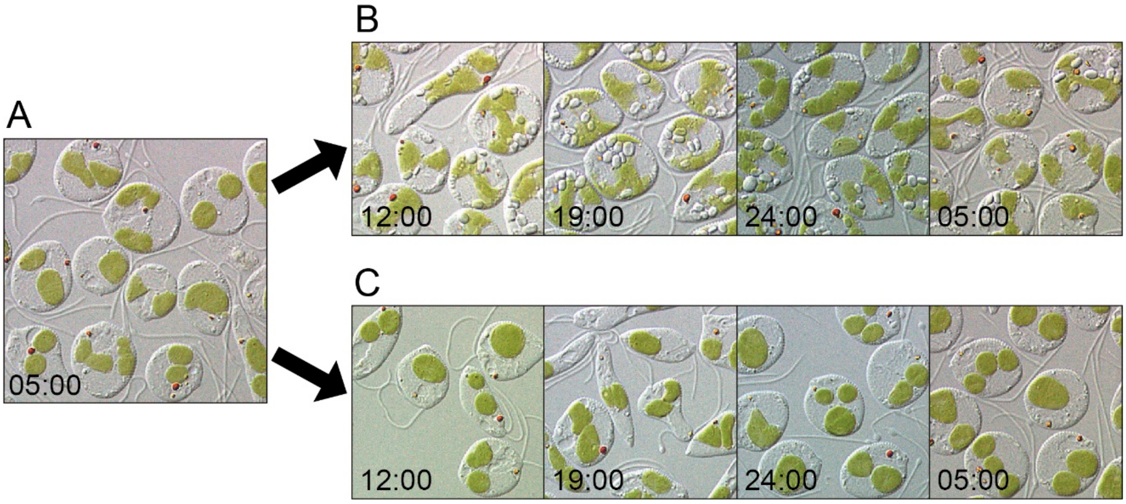

**Fig. S8.** Diverse origins of the kleptoplast-targeted sequences in *Rapaza viridis*. Maximum likelihood trees were estimated with 1,000 rapid bootstrapping replicates in IQ-TREE. Photosynthetic Euglenids are written in bold, and *Rapaza* is also written in purple. Black circles (•) denote bootstrap support >95; support values below 50 are not shown. A: Phylogeny of *ftsH* suggest a green algal origin of plastid-targeted genes in all euglenophytes. B: Phylogeny of transketolase suggesting a gene transfer from the red algae-derived complex plastids to Euglenophyceae.

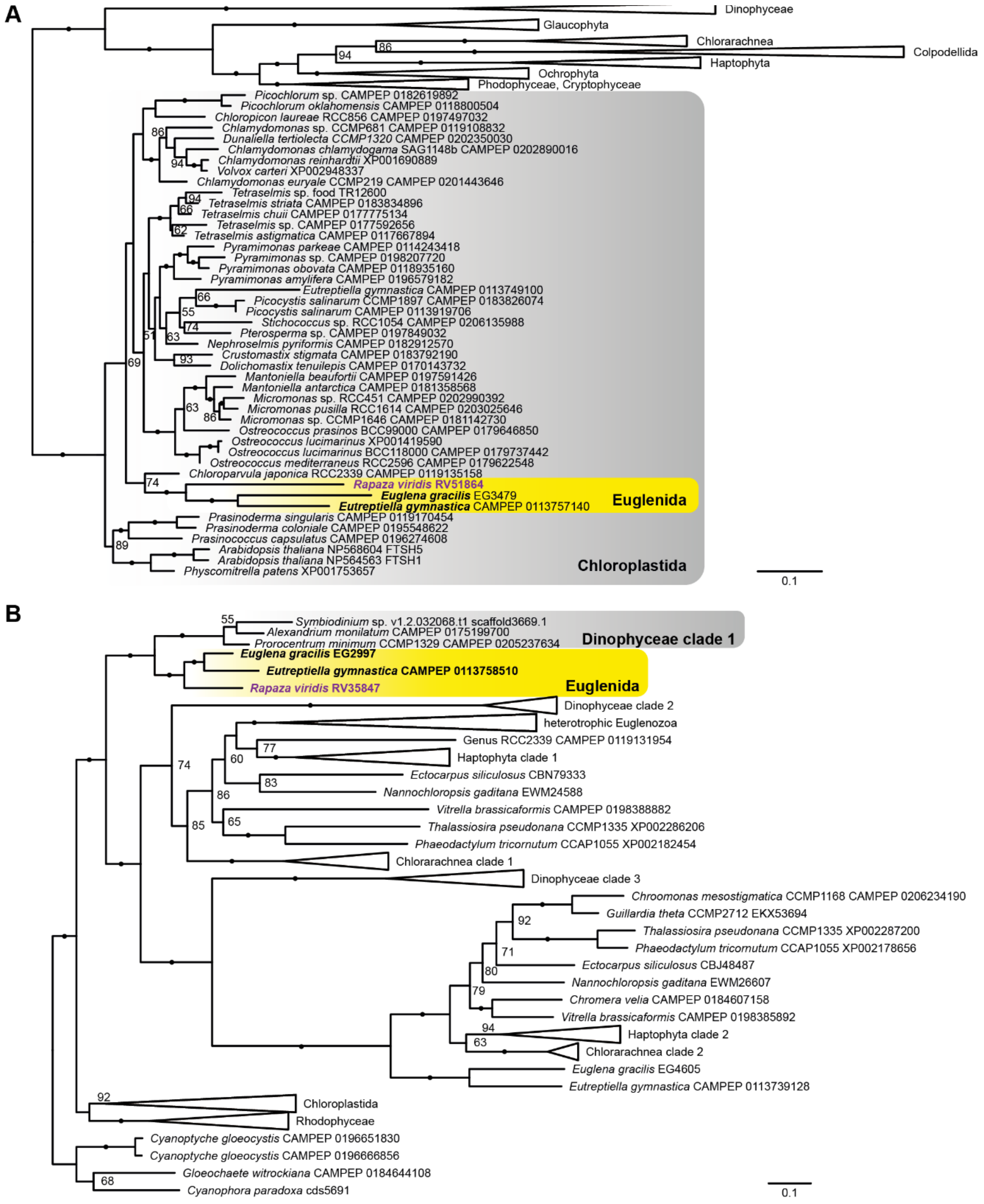

**Table S1.** $\delta^{13}C$ values of polysaccharide grains purified from the *Rapaza viridis* cells incubated in Daigo-IMK medium with and without the addition of trace amounts of $^{13}C$-labelled bicarbonate; and the $\delta^{13}C$ value of the labeled medium. The labeling experiment clearly indicated the uptake and fixation of inorganic carbon by *R. viridis*. The results indicate that 48% of the carbon constituting the polysaccharide grains formed during the incubation period was derived from inorganic carbon taken up extracellularly from the medium.

| Sample | $\delta^{13}C$ value |
|---|---|
| Purified polysaccharide grains from the $^{13}C$-labelled experiment | 33 |
| Purified polysaccharide grains from control experiment | -22 |
| Residue of the $^{13}C$-labelled medium after dry | 91 |

**Table S2 (separate file).** 274 candidates for plastd-targeted proteins in the predicted *Rapaza viridis* proteome.The translated transcriptome of *R. viridis* have been annotated with BLAST at the E. gracilis plastid proteome and UniProt database. The potential membrane-spanning regions were identified with the hidden-Markov-model-based program TMHMM; the sequences with at least one transmembrane domain were kept resulting in 274 candidates.

**Table S3 (separate file).** 63 candidates for plastd-targeted proteins in the predicted *Rhabdomonas costata* proteome.The translated transcriptome of *R. costata* have been annotated with BLAST at the *E. gracilis* plastid proteome and UniProt database. The potential membrane-spanning regions were identified with the hidden-Markov-model-based program TMHMM; the sequences with at least one transmembrane domain were kept resulting in 63 candidates.

**Movie S1 (separate file).** *Rapaza viridis* preying on *Tetraselmis* sp., movie starting immediately after addition of the *Tetraselmis* cells to the *R. viridis* culture. Within minutes, *R. viridis* ingested the whole *Tetraselmis* cell by phagocytosis.

1.
